# An experimental evaluation of drug-induced mutational meltdown as an antiviral treatment strategy

**DOI:** 10.1101/048934

**Authors:** Claudia Bank, Nicholas Renzette, Ping Liu, Sebastian Matuszewski, Hyunjin Shim, Matthieu Foll, Daniel N. A. Bolon, Konstantin B. Zeldovich, Timothy F. Kowalik, Robert W. Finberg, Jennifer P. Wang, Jeffrey D. Jensen

## Abstract

The rapid evolution of drug resistance remains a critical public health concern. The treatment of influenza A virus (IAV) has proven particularly challenging, due to the ability of the virus to develop resistance against current antivirals and vaccines. Here we evaluate a novel antiviral drug therapy, favipiravir, for which the mechanism of action in IAV involves an interaction with the viral RNA-dependent RNA polymerase resulting in an effective increase in the viral mutation rate. We utilized an experimental evolution framework, combined with novel population genetic method development for inference from time-sampled data, in order to evaluate the effectiveness of favipiravir against IAV. Evaluating whole genome polymorphism data across fifteen time points under multiple drug concentrations and in controls, we present the first evidence for the ability of viral populations to effectively adapt to low concentrations of favipiravir. In contrast, under high concentrations, we observe population extinction, indicative of mutational meltdown. We discuss the observed dynamics with respect to the evolutionary forces at play and emphasize the utility of evolutionary theory to inform drug development.

## INTRODUCTION

The increasing availability of time-sampled data, and of statistical inference approaches based on such data, is considerably expanding the repertoire of population genetics (reviewed in Bank *et al.* 2014). Time-sampled data are often considered in association with the growing field of ancient DNA and have long been a key feature in the analysis of both experimental and clinical data. Indeed, one of the most important applications and challenges today is how to best utilize such data in order to characterize the evolution of human pathogens, perhaps most specifically, how to quantify (and ultimately combat) the ability of viral populations to develop resistance to given treatment strategies.

Influenza A virus (IAV) is of long-term public health interest given its scope (with approximately 36,000 deaths annually in the United States alone; Thompson *et al.* 2003) and the rapid evolution of resistance against common therapeutics. The evolution of resistance is ultimately a process dictated by the nature of the interaction between drug, virus, and host. For example, the most widely administered drug for combating IAV, oseltamivir, was initially designed as a competitive inhibitor of neuraminidase based on structural information of the active site (Moscona 2005; Colins *et al.* 2008). Though it was widely believed that resistance to oseltamivir would be clinically unimportant given the associated high fitness cost (Ives *et al.* 2002), a particular resistance mutation neuraminidase H275Y nonetheless spread rapidly (Gubareva *et al.* 2001; Moscona 2009; Ghedin *et al.* 2012) - likely due to the presence of accompanying compensatory mutations (Bloom *et al.* 2010; Bouvier *et al.* 2012; Ginting *et al.* 2012). Hence, in the case of oseltamivir, a single mutation near the viral neuraminidase active site is sufficient to attenuate drug binding and thereby cause resistance. Thus, alternative classes of drugs for which resistance evolution is less easily achieved have garnered interest.

One promising drug is favipiravir, for which the mechanism of action is related to the selective inhibition of viral RNA-dependent RNA polymerase (RdRp) and the decrease of polymerase fidelity, which results in an increase of the genome-wide mutation rate in IAV (Baranovich *et al.* 2013; Furuta *et al.* 2013). To our knowledge, no study to date has found evidence for successful resistance evolution against favipiravir. This may suggest either that resistance is complex to achieve given that the drug effectively targets the entire genome (and thus no simple genetic solution for resistance exists), or that its effect is so strong that viral populations are always driven to extinction prior to the appearance of resistance mutations.

The field of evolutionary theory has studied the effects of increasing mutation rates on asexual populations for decades, and several processes may be invoked that lead to the extinction of a population owing to an artificially increased mutation rate. All are based on the fact that the majority of new fitness-affecting mutations are deleterious.

First, Muller’ s ratchet is a process that describes the decline of fitness and size of a non-recombining population, owing to the periodic loss of the most fit genotype (Muller 1964; Felsenstein 1974). Theoretical work has shown that the speed of the ratchet is determined by the effective population size, mutation rate, deleterious selection coefficient, and the size of the least-loaded (i.e., most fit) class at mutation-selection equilibrium (Haigh 1978, and see Gordo and Charlesworth 2000). Although the ratchet initially leads to a linear accumulation of mutations while census population size remains constant, it may ultimately lead to the rapid extinction of the population due to what is referred to as “mutational meltdown” (Lynch *et al.* 1990; Lynch *et al.* 1993). Under high mutation rate conditions, the loss of the least-loaded class may be primarily driven by mutation rather than genetic drift (Lynch *et al.* 1993). We argue that the process of mutational meltdown is essentially similar to “lethal mutagenesis”, a term that has subsequently been promoted in the area of virology (Bull *et al.*, 2007; Wylie & Shakhnovich, 2012). Both occur in finite populations under high mutation rates, and are characterized by the linear accumulation of mutations until the mean viability of individuals is too low to maintain the carrying capacity, at which point both population size and fitness deteriorate rapidly, leading to extinction. They differ in two main ways: a primary focus on effective population size (mutational meltdown) versus census population size (lethal mutagenesis), and in the assumption of strong (mutational meltdown) and weak (lethal mutagenesis) genetic drift. However, extinction is eventually driven by mutation load in both, and the models are thus difficult to distinguish empirically.

Second, although beneficial mutations (e.g., mutations conferring drug resistance or other relative growth advantages) will also become more frequent under increased mutation rates, these are likely to occur in individuals that also carry deleterious mutations, which may prevent or slow their spread in the population in the absence of recombination. This effect, termed “weak-selection Hill-Robertson interference” (WSHRI) (Hill and Robertson 1966; McVean 2000), thus may also serve to slow the evolution of resistance.

Finally, the extinction of populations under high mutation rate conditions has been frequently discussed in the area of virus evolution with regard to the concept of error catastrophe (Eigen 1971; Eigen 2002; Holmes 2003). In principle, the concept of error catastrophe is similar to the theories of mutational meltdown and lethal mutagenesis, and large parts of the theory are equivalent (Wilke 2005). However, the causal mechanism of extinction due to error catastrophe is specifically mutation accumulation beyond the “error threshold” above which the evolutionary dynamics destabilize and the genome is unable to maintain the required information. The error threshold is a sharp limit, and the occurrence of error catastrophe is not limited to finite populations.

Here we present data from *in vitro* selection experiments in which populations of IAV were evolved in the presence or absence of favipiravir treatment and compare these to our previous, similar experiment using oseltamivir (Foll *et al.* 2014; Renzette *et al.* 2014). Results indeed demonstrate that favipiravir induces mutational meltdown, with experimental populations exposed to escalating drug concentrations eventually becoming extinct. This result is in stark contrast to the populations that evolved in the presence of escalating amounts of oseltamivir, in which resistance mutations arose and fixed quickly after the introduction of drug pressure. We also evaluate different concentrations of favipiravir treatment, quantifying the extent necessary to induce this effect, and we provide the first evidence for potential adaptation to favipiravir under a constant low drug concentration.

As such, this experimental set-up allowed us to directly study the evolutionary dynamics of IAV in different drug environments. In doing so, we were able to quantify the adaptive process with respect to potential beneficial
mutations in response to different conditions and discuss the potential for evolutionary rescue (i.e., the process by which a population can escape extinction in a novel environment through rapid adaptation; Alexander *et al.* 2014). Thus, this work highlights experimental scenarios of clinical relevance of both successful and failed evolutionary rescue, and allows us to observe the dynamics of mutational meltdown in action. Our results demonstrate the promise of drug-induced mutational meltdown as a means for combating viral populations, and for favipiravir as an effective strategy against IAV in particular. Yet, our findings also raise concerns that proper drug dosage is essential for effective treatment. By discussing our results with respect to concepts from evolutionary theory, we outline prospects for the better prediction of the evolutionary response of pathogens to drug pressure.

## MATERIALS & METHODS

### Cells, virus stocks, and chemicals

Madin-Darby canine kidney (MDCK) cells were obtained from American Type Culture Collection (Manassas, VA) and propagated in Eagle’ s minimal essential medium (MEM) with 10% fetal bovine serum (FBS; Hyclone, Logan, UT) and 2 mM penicillin/streptomycin. Influenza virus A/Brisbane/59/2007 (H1N1), grown in the chicken egg allantoic fluid, was obtained through the NIH Biodefense and Emerging Infections Research Resources Repository, NIAID, NIH (NR-12282; lot 58550257). Favipiravir was obtained from FUJIFILM Pharmaceutical USA, Inc.

### Viral titer determination by plaque assay

Viruses were quantified on MDCK cells to determine infectious titer (plaque forming units per ml, or PFU/ml) as previously described (Hendricks *et al.* 2013). In brief, six 10-fold serial dilutions were performed on the viral samples followed by 1 h of binding at 37°C on confluent MDCK cells in 12-well plates. After washing off unbound virus with phosphate buffered saline (PBS), the cells were overlaid with agar (0.5%) in DMEM-F12 supplemented with penicillin/streptomycin, L-glutamine, bovine serum albumin, HEPES, sodium bicarbonate, and 20 μg/ml acetylated trypsin (Sigma, St. Louis, MO). After the agar solidified, the plates were incubated for ~48 h at 37 °C. Cells were fixed and stained with primary antibody anti-H1 (MAB8261, Millipore, Billerica, MA). Plaques were visualized with anti-mouse horseradish peroxidase-conjugated secondary antibody (BD Biosciences, San Jose, CA) and developed with peroxidase substrate kit (Vector Laboratories, Burlingame, CA).

### Determination of favipiravir ED_50_

The 50% effective dose (ED_50_) value was defined as the concentration of drug reducing plaque number to 50% of no drug control. In brief, the ED_50_ was determined by seeding 2.5 × 10^5^ MDCK cells in each well of a 24-well plate and incubated overnight at 37 °C, 5% CO2. Virus was added to cells at an MOI of 0.01 in 100 μl of IAV growth medium [EMEM/10% FBS with 2 mM penicillin/streptomycin, 7.5% bovine serum albumin, and 1 μg/ml TPCK-treated-trypsin (Sigma)] plus favipiravir (0, 0.1, 0.3, 1, 3, or 10 μM). After incubation at 37 °C for 1 h, cells were washed once with PBS; 500 μl of IAV growth medium with the appropriate concentration of favipiravir was added and cells were again incubated at 37 °C for several days. Supernatants were collected when > 50% cytopathic effect (CPE) was achieved for at least one drug concentration. Supernatants were centrifuged for 15 min at 300 *x g* at 4 °C and stored at −80° C. The viral titer for each sample was determined by plaque assay.

### Quantitative PCR

Viral RNA was extracted from supernatants using the QIAamp Viral RNA Mini kit (Qiagen), then reverse transcribed with the High Capacity cDNA Reverse Transcriptase Kit (ThermoFisher Scientific). Viral copies were quantified using M1 forward primer AAGACCAATCCTGTCACCTCTGA, reverse primer CAAAGCGTCTACGCTGCAGTCC, and probe TTTGTGTTCACGCTCACCGT for 40 cycles using the Eppendorf Mastercycler ep Realplex program.

### Viral culture

Viruses were serially passaged in MDCK cells (2.5 × 10^5^ cells per well). For the first experiment (favi1, constA, withdrawalA, and its controls), the multiplicity of infection (MOI) was generally fixed at 0.01 for each passage. The MOI was occasionally adjusted to accommodate for the available volume/titer of virus, including for escalating favipiravir (MOI 0.001 for P15) and for constant favipiravir/withdrawal of favipiravir (MOI 0.001 for P11, MOI 0.005 for P15).

For the second experiment (favi2 and its control), virus was harvested as the culture reached ~50% CPE, and the cell-free virus processed for sequencing. The variable MOI was a result of continuously passaging the viral populations. At each passage, a range of virus was used to initiate the next round of infection in various wells. The sample that generated 50% CPE with the lowest input of virus was used to continue the trajectory. Virus titers were determined at the conclusion of the experiments. The average MOI was subsequently calculated as 0.02 ± 0.009 (S.E.M.).

Samples were harvested both in the presence and absence of escalating amounts of favipiravir. In the first passage with favipiravir, the drug concentration was 2X the ED_50_ of 1 μM (2 μM), consistent with reports of seasonal strains of IAV, ranging from 0.45 to 5.99 μM, (Sleeman *et al.* 2010). The concentration was doubled for each subsequent passage as long as > 50% CPE was present. If < 50% CPE was present, the concentration of favipiravir was escalated at a slower rate.

### High-throughput sequencing

We developed a high-throughput sample processing work flow, carried out in 96-well format, including RNA purification, reverse transcription, whole genome PCR, followed by DNA barcoding and library preparation. See Renzette *et al.* (2014) for details.

### Bioinformatics analysis

Short reads from the Illumina or IonTorrent platform (see Supplementary Table 1) were filtered for quality scores > 20 throughout the read and aligned to the strain’ s reference genome using BLAST. Over 95% of the selected reads could be mapped to the IAV reference genome obtained from GenBank (accessions CY030232, CY031391, CY058484-CY058486, CY058488-CY058489, CY058491). Only alignments longer than 80 nucleotides were retained. The median sequencing depth was 10,226. Amino acid frequencies were calculated after aligning translated reads to the corresponding positions in the reference proteins. We confirmed that nucleotide and amino acid frequencies were identical between passages. Unfolded SNP frequencies were generated using the IAV reference genome and were used for the population genetics analyses. The sequencing datasets generated in this study are available at http://bib.umassmed.edu/influenza. See Renzette *et al.* (2014) for further details.

### Sequencing error analysis

See Renzette *et al.* (2014) for further details. Because segment ends are known to contain repetitive regions that are difficult to map, the first and last 30 sites of each segment were excluded from all analyses.

### Population dynamics

We calculated absolute growth rates for all treatments based on the initial and final population size (obtained directly before and after virus passaging) as given by the MOI and the PFU/ml, respectively (see Supplementary Table 1). Note that the latter implicitly assumes that each formed plaque represents a single infective particle. However, given the generally low MOI this assumption should hold true for our data. Following Foll *et al.* (2014), we assumed that for all treatments the population grew for 13 generations at constant rate *r* per passage, yielding

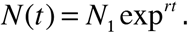

Note that after rearranging and solving for *r*, the corresponding absolute growth rate is an estimate for the Malthusian parameter or the intrinsic rate of increase Chevin (2010).

Selection intensity *s* between different treatments was quantified by calculating *s = r_treatment_ - r_control_*. We then performed a linear regression via ordinary least squares of the growth rates *r* and selection coefficients against time (given by passage number) or drug concentration. To assess whether there was a significant linear relationship between these entities (e.g., growth rate versus initial population size; see Fig 3A), we performed an ordinary t-test under the null hypotheses of no significant relationship between the independent (i.e., drug concentration, passage or MOI) and dependent variable (i.e., absolute and relative growth rate), corresponding to a zero slope. To test whether two regression slopes were significantly different from one another we again performed an ordinary t-test under the null hypotheses that both slopes are equal.

### WFABC analysis

We utilized the WFABC software (Foll *et al.* 2014; Foll *et al.* 2015) to estimate the effective population size *N_e_* and selection coefficients from allele-frequency trajectories. Given the sequencing error estimates of up to 1% (see Supplementary Figure 1), we randomly down-sampled sites with coverage above 100 to a sample size of 100. At each site we kept the counts of the ancestral allele and the most frequent derived allele, hence we consider only diallelic SNPs (Foll *et al.* 2014). For WFABC we kept only trajectories that had a derived allele count of at least 3 at one passage throughout the experiment, corresponding to a frequency of >2.5% in the population. We used a uniform prior probability of - 0. 5 < *s* < 0.5 This resulted in the following command line for the wfabc_2 selection estimation step: ./wfabc_2-ploidy 1-min_s-0.5-max_s 0.5-min_freq 0.025 FILE

For the “fork” data sets, we ran WFABC both for passages 9-17, (denoted by “constA” etc. in Supplementary Table 2) and for the full time series of passages 3-17 (denoted by “favi1-constA” etc in Supplementary Table 2). For the favi1 data set, we used both the full range of passages 3-15 (favi1-long), and a reduced data set excluding the last passage (favi1-short). Consistent with Foll *et al.* (2014), candidate trajectories under positive selection were selected to be those with a posterior probability of s<0 less than 0.5%. Passage 12 in constA and constB, and passage 11 in favi2 and favi2-control were excluded from the analysis because of low coverage.

### Hierarchical clustering analysis

Our hierarchical clustering analysis (Figure 4) is based on the squared Euclidean distance between allele-frequency trajectories of the candidate mutations identified by WFABC using Ward’ s minimum variance criterion (Ward Jr. 1963). Furthermore, for all WFABC candidates we calculated the correlations between allele trajectories - starting from the point in time where their frequency was above the level of the sequencing error (i.e., above 1%) - and the absolute and relative growth rate estimates (Supplementary Table 2).

### CP-WFABC analysis

The CP-WFABC approach is an extension of WFABC, and considers models of changing *N_es_* from time-sampled polymorphism data (Shim *et al.*, 2016). The method uses the variance in allele frequencies between two consecutive sampling time points defined as *Fs′*, an unbiased estimator of *N_e_*, to measure selection strength (Jorde and Ryman 2007):

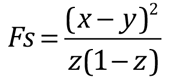

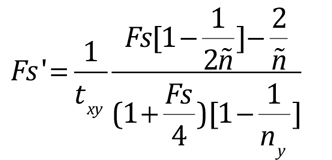
where *x* and *y* are the allele frequencies at two time points separated by *t_xy_* generations, *z = (x+y)/2*, and *ñ* is the harmonic mean of the sample sizes *n_x_* and *n_y_* at those time points. Additionally, the method adapts a procedure from Change-Point analysis in time-serial data to detect a change point of selection along the allele trajectory. More specifically, a statistic from the cumulative sum control chart (CUSUM) developed by Page (1954) is integrated into the ABC method to characterize the time-sampled trajectory of an allele:

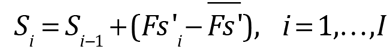
where the index *i* refers to the *i*th time point, *S*_0_ = 0, and 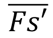 corresponds to the mean of the *Fs’* over all pairs of consecutive time points. The change point *S_CP_* is the sampling time point with the maximal absolute value of *S_m_*, which is the maximal accumulation of difference in *Fs’* from its average value in the time-sampled trajectory:

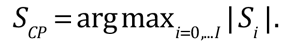

Using these two summary statistics (*Fs*’ and *S_cp_*), CP-WFABC jointly estimates the temporal position of a change point as well as the strength of both corresponding selection coefficients (and dominance for diploid cases) from an allele trajectory. Furthermore, CP-WFABC separates allele trajectories characterized by a single selection coefficient from those with changing intensity via ABC model choice (see below).

The same data sets and ascertainment conditions were used as for the WFABC analysis described above. For each trajectory of interest, we tested the null model M_0_ (i.e., a single selection coefficient) and the alternative model M_1_ (i.e., a changing selection coefficient), with the parameters of interest being (*s*) and *(s1, s2, CP)*, respectively. The uniform prior ranges for the selection coefficients were set as [-1,1]. The prior here was chosen to be wider than for the traditional WFABC analysis described above since CP-WFABC allows for the detection of strongly deleterious effects after the change point. We chose a uniform prior for the change point ranging from the second generation to the second-to-last generation. 1×10^6^ simulated datasets were generated using the Wright-Fisher model for M_0_ and M_1_, while keeping the other input values, such as the effective population size estimated from WFABC, the number of generations, the sampling time points, and the sample sizes identical for each observed trajectory. The best 1×10^3^ simulated trajectories were drawn from the combined M0 and M1 sets, determined by the smallest Euclidean distance of the aforementioned summary statistics between the simulated trajectory and the observed trajectory. The approximate posterior densities of the parameters of interest for M_0_ and M_1_ were built using the respective subsets of these chosen simulations, as in the algorithm described in Beaumont *et al.* (2002). Additionally, the ABC model choice was constructed to identify the SNP trajectories with changing selection intensity, with the relative probability of M_1_ over M_0_ as the model’ s posterior ratio and as the Bayes factor *B_1,0_* (Sunnåker *et al*, 2013):

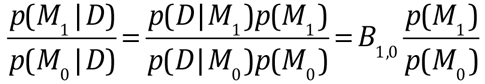

when the model prior *p*(M_0_) is equal to *p*(M_1_). For a small haploid population, as in the case for IAV in this experimental setup (i.e., *N_e_* on the order of hundreds), the Bayes factor *B_1,0_* must exceed 3.7 in order to achieve sufficient support for the alternative model M_1_ with a significance level of 1% (see Supplementary Figure 9).

### Population size estimates

In order to estimate the effective population size between any two time points, we used the *Fs’* statistic introduced above. For each pair of time points and each site in the genome, we (hypergeometrically) sampled min(100, COVERAGE) reads from the focal data set, and included these values in our calculation if one of the observed frequencies was >2.5%. We then computed the estimated effective population size for this pair of time points as 1/mean(*Fs’*).

### Accumulation of mutations

To infer the average number of mutations per individual accumulating over time, which is not directly observed due to the lack of haplotype information in the data, we took the sum over the derived allele frequencies at all sites with coverage greater than 100, if the frequency was above the sequencing error threshold of 1%. We then extrapolated this value to the whole genome.

## RESULTS & DISCUSSION

### The experimental setup

We analyzed the evolution of IAV across nine experimental populations exposed to different drug conditions as illustrated in Figure 1. As explained in detail in the Materials & Methods section, the virus (H1N1) was serially passaged on Madin-Darby Canine Kidney (MDCK) cell culture, and we assumed each passage to be an average of 13 viral generations (see Foll *et al.* 2014). Each treatment population (left half of Figure 1) was accompanied by a control population passaged in the absence of the drug (right half of Figure 1) to account for environmental fluctuations, minor variation in MDCK cell culture conditions, and multiplicity of infection (MOI) (see Materials & Methods). The stock viral populations originated from passage 3 of an earlier experiment described in Foll *et al.* (2014) and Renzette *et al.* (2014). At this point, the population was split into four subsets (hereafter referred to with the respective abbreviations): two replicates with an increasing concentration of favipiravir (favi1, favi2) and their parallel controls that were not exposed to the drug (favi1-control, favi2-control). After passage 9, three additional populations were created from favi1, paralleled by two additional populations from its accompanying favi1-control population; these populations are hereafter referred to as “forks”. These forks consisted of two populations originating from favi1 exposed to a constant concentration of favipiravir (constA and constB) and their accompanying controls without drug originating from favi1-control (constA-control, constB-control), as well as a fifth fork in which favipiravir was withdrawn, originating from favi1 (withdrawalA; i. e., drug pressure was completely halted after passage 9). A summary of MOIs, drug concentrations, and other specifics are provided as Supplementary Table 1.

**Figure 1.**
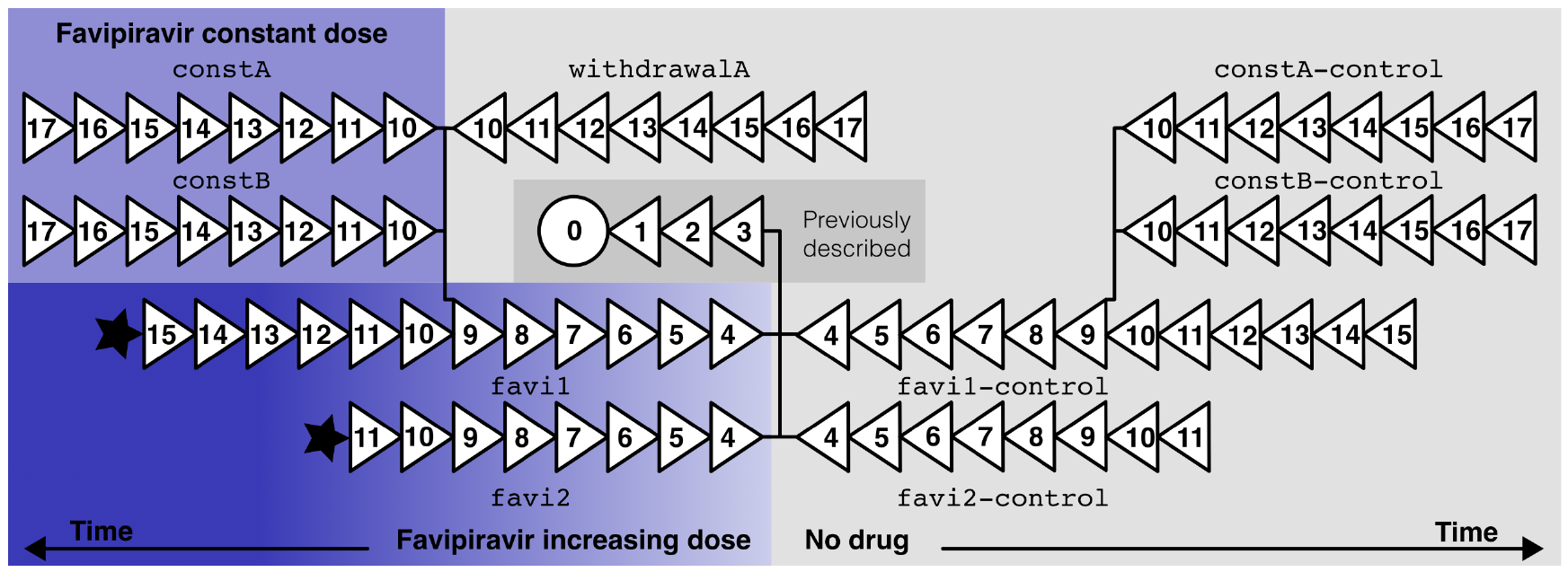
Experimental evolution of IAV with and without favipiravir. Each triangle represents an experimental passage, at the end of which whole-genome sequencing was performed. Black stars indicate extinction of the viral population. Rows (excluding the dark gray rectangle) represent parallel sets of treatments and control experiments that were processed and sequenced in the same sequencing lane (ensuring similar effects of cell culture and sequencing protocol). Labels represent the names of the populations subsequently used in the manuscript. See Supplementary Table 1 for information on MOI and drug concentrations.

The favi1 and favi1-control populations were continued until passage 15, after which time too few virions were recovered from favi1 to continue the experiment. Similarly, favi2 and favi2-control were discontinued after passage 11 - these events are hereafter referred to as extinctions.

In the following, we quantify and discuss the observed patterns from an evolutionary perspective, with a focus on dissecting the process that leads to extinction of the viral population in favi1 and favi2, and discuss the potential for the evolution of resistance against favipiravir.

### Evidence for increased mutation rate under favipiravir treatment

Favipiravir affects IAV by increasing the mutation rate above sustainable levels, leading to an accumulation of deleterious mutations and the eventual extinction of the population (Furuta *et al.* 2013). We sought to validate the mutation-rate increasing effect of favipiravir on IAV that was previously described by Baranovich *et al.* (2013). However, the quantification of mutation rates from genomic data is obscured by sequencing error, which induces a (false) baseline of observed variation. To investigate the effect of favipiravir on the mutation rate while accounting for this complication, we studied the number of segregating mutations above two frequency thresholds, *f* of 0.1% and 1%. Whereas the number of segregating sites above *f* = 0.1% is likely strongly confounded by sequencing errors and expected to vary depending on sequencing depth and quality, *f* = 1% is expected to be above the sequencing error threshold (see Supplementary Figure 1), but few mutations will rise to such high frequencies. We estimated the number of segregating sites in the genome by discounting all sites with coverage lower than 1/*f* (which for most passages comprised the majority if not the entirety of the genome), and, if necessary, extrapolated this value to the whole genome. The results are illustrated in Figure 2 and Supplementary Figure 2.

**Figure 2.**
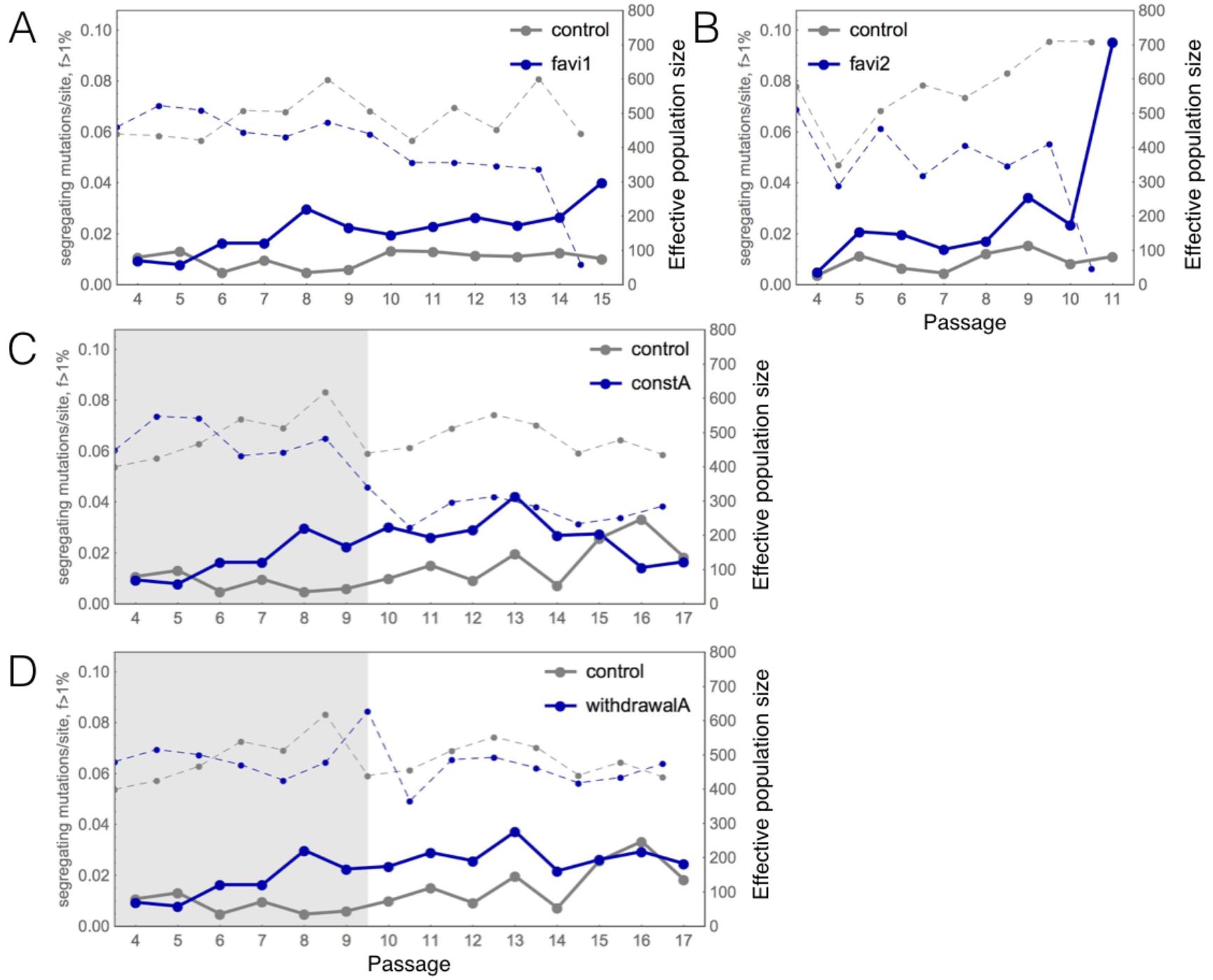
The number of mutations per site in the genome that segregated above 1% across time and populations (left y axis), with data points connected by solid lines. In the background, estimates of the effective population size between any two passages are displayed (right y axis, data points connected by dashed lines). (A-B) We observed a greater number of segregating mutations in the favipiravir treatments favi1 and favi2 as compared with control populations, consistent with the proposed mechanism of an increase in mutation rate. In the favi2 population, a particularly steep increase in the number of segregating mutations occurred immediately prior to extinction (B). Also the constA (C) and withdrawalA (D) treatments showed an increased number of segregating mutations as compared with their control experiments, but the accumulation of mutations comes to a halt in withdrawalA, and appears to slowly recover in constA. The effective population size tends to be lower in the treatment populations (blue) than in the controls (gray). Gray shading indicates that this part of the figure represents favi1 increasing data, before forks were created. Note that although passages 4 to 9 in panels A, C, and D stem from the same data, estimated population sizes differ slightly due to the sampling procedure that was individually performed to calculate the *Fs’* statistic.

For the lower threshold *f* = 0.1%, we observed a strong correlation between the number of segregating mutations in any two parallel experiments (see Supplementary Figure 2). This is expected if sequencing error is prevalent at this frequency threshold. Hence, differences in the number of segregating mutations in this frequency range reflect differences in coverage and sequencing quality across passages and experiments.

For the high threshold *f* = 1%, we observed an increase in the number of segregating mutations with time in the favi1 and favi2 populations, consistent with an increase in the mutation rate due to favipiravir treatment (see Figure 2A–B); we discuss alternative explanations, including different or changing effective population sizes, or selected mutations, below. In every population associated with favipiravir treatment (favi1, favi2, constA, constB, withdrawalA), the average number of segregating mutations was higher than in any of the control treatments. Whereas in favi1 the increased number of mutations was clearly visible by passage 8, favi2 appeared to be affected even at very low concentrations. These differences between favi1 and favi2 were likely due to the constant and larger bottleneck sizes between passages in favi1 as compared with favi2 (see Materials & Methods, Supplementary Table 1). The observed increase in the number of segregating mutations was roughly 2.5-fold (excluding the last passage of favi2) which is in agreement with previous estimates of the relative increase in mutation rate under favipiravir treatment (Baranovich *et al.* 2013).

For constA, constB and withdrawalA (Figure 2 C–D, Supplementary Figure 2A) the number of segregating mutations remained greater than that of the control for the most part, but we observed no further increase as in favi1 and favi2. As the experiment ended at passage 17, the potential recovery of the constA population that is indicated in Figure 2C (but see below) cannot be evaluated.

### Monitoring the effective population size throughout the experiment

One alternative explanation for the increasing number of segregating mutations could be a changing effective population size through time between treatments. If most new mutations are deleterious, larger effective population sizes may result in a lower number of segregating mutations due to an increasing efficacy of selection. We estimated the effective population sizes between any two passages based on the *Fs’* statistic postulated by Jorde & Ryman (2007) (see Materials and Methods). *Fs’* uses the variance of allele frequencies between two time points to evaluate the strength of genetic drift in the population, and should therefore not be affected by an increased mutation rate (but see below for a discussion of the assumption of independence of sites). The estimated effective population sizes are displayed in the background of Figure 2.

Across all populations and passages, the estimated effective population size (i.e., 1/mean(*Fs’*), with the mean being taken over the *Fs’* values between each pair of consecutive time points; see Materials and Methods) ranged from 48 between passage 10 and 11 in the favi2 population to 823 between passage 10 and 11 in the constB-control, which is consistent with previous estimates (Foll *et al.* 2014). The average estimated effective population sizes correlate well with the estimated global *N_e_* from WFABC (see Supplementary Figure 3). Population sizes were slightly but consistently larger in the control populations, and in the favi1 and favi2 environments we observe a steep decline in the effective population size immediately prior to extinction (see Figure 2A–B). Interestingly, upon withdrawal of the drug pressure, the population size recovered and continued at a similar size as its control despite a greater number of segregating mutations (see Figure 2D). Conversely, in the constA population, the effective population size remained consistently lower than both its control and the withdrawal population throughout the entire course of the experiment (see Figure 2C).

### Population dynamics across drug conditions

In contrast to data from natural populations, the experimental setup of serial passaging of the virus in cell culture allowed us to control and monitor the population dynamics of the virus, and thus to directly assess the number of virions introduced to the cell culture (reflected in the multiplicity of infection [MOI]), and the number of virions emerging at the end of each passage (output virions, specifically plaque-forming units [PFU]; see Supplementary Table 1). We used these figures to estimate absolute growth rates of the viral population during each passage under the assumption of exponential growth (see Figure 3, Supplementary Figure 4, and Materials and Methods).

**Figure 3.**
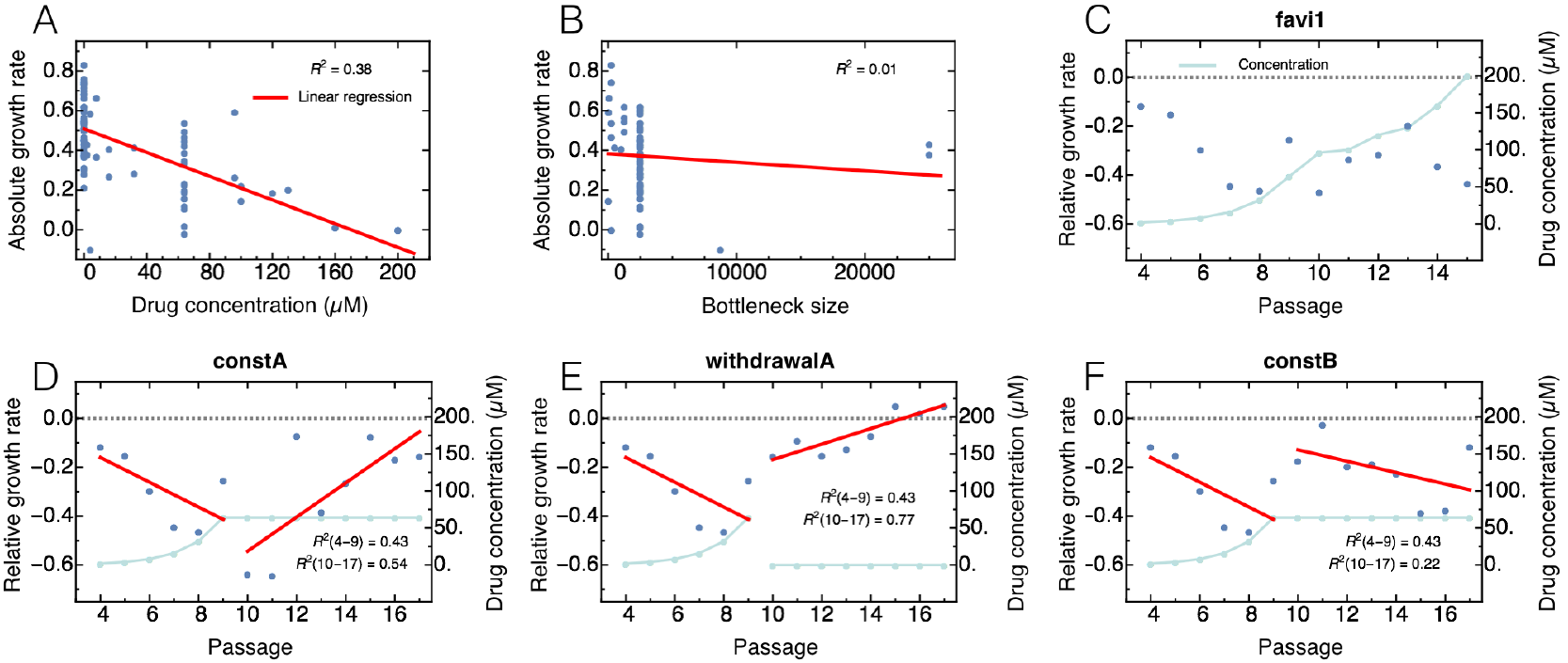
Changes in absolute and relative growth rates of the virus throughout the experiment. Panel A: Absolute growth rate showed a strong negative correlation with imposed drug concentration, providing further evidence for the effectiveness of the treatment. Panel B: No correlation was observed between the initial population size in each passage and the absolute growth rate. Panel C: In favi1, relative growth rates (compared with parallel control) were consistently negative. Panels D-F: Relative growth rates of additional treatment strategies across passages. Whereas the growth rate decreased in the first part of the experiment (favi1, passage 4-9) as drug concentration (blue line) increased, the constA (panel D) and withdrawal (panel E) populations showed signs of recovery upon the change of treatment.

As a further assessment of the effectiveness of the drug treatment, we studied the relationship between the drug concentrations throughout all populations and the absolute growth rates, and reassuringly observed a highly significant negative correlation (Figure 3A; R^2^=0.38 p<10^-9^). While there appears to be no correlation globally between the bottleneck size and the growth rate (Figure 3B; R^2^=0.01 p=0.26), indicating that the imposed population dynamics do not significantly impact the rate of growth, a small negative effect of bottleneck size on the estimated growth rates cannot be ruled out when only considering bottleneck sizes of 5000 and smaller (R^2^=0.11 p=0.009; see also Supplementary Table 3).

To study the effect of different drug conditions on the growth rates, we computed relative rates as the difference between the growth rate of the treatment and its parallel control, *s* = *r_treatment_ - r_control_*. This accounts for effects due to the MDCK cell environment and the size of the imposed passaging bottlenecks, which were generally kept identical between parallel experiments (see Supplementary Table 1).

We observed consistently negative relative growth rates in the favi1 and favi2 populations (Figure 3C and Supplementary Figure 4), likely due to the challenging effect of the drug on the population. It is important to note differences in the experimental procedure in the favi2 population and its parallel control from that of all other populations (see also Materials & Methods). In general, bottleneck sizes (i.e., MOIs) were kept constant and relatively large, and new passages were seeded from a random sample of the previous population. However, the bottleneck sizes in favi2 were smaller and varied greatly, and new passages were seeded based on the fastest-growing amongst several samples. Therefore, slightly different patterns between the favi1 and favi2 populations are expected (such as the higher absolute growth rates in favi2 (see Supplementary Figure 4)). Furthermore, the lower MOIs in favi2 may have resulted in a lower probability of co-infection of cells, and thereby in a lower chance of virus segment reassortment which, analogous to recombination, could help to bring together co-adapted segments or to purge deleterious mutations from the viral genome; a potential consequence is a more rapid extinction of the favi2 population.

Interestingly, in the constA population (Figure 3D), the initially negative relative growth rate appeared to gradually recover under constant drug pressure, ultimately approaching 0 (i.e., similar to the growth rate of the parallel control). The slope of a linear regression from passage 10 to 17 is positive (p=0.039). This provides the first line of evidence that the constA population may be acquiring resistance under low-concentration favipiravir conditions. A similar pattern of recovery (increasing slope of linear regression, p=0.004) was observed in the withdrawalA population (Figure 3E). Finally, the constB population did not show signs of recovery (p=0.24), but rather maintained a constant negative relative growth rate throughout treatment (Figure 3F).

### Identification of putatively selected mutations

The whole-genome SNP data obtained at the end of each passage in all populations provided us with allele-frequency trajectories through time for every site in the genome. These trajectories contain information on the effective population size and thus on the magnitude of genetic drift, and on the selection coefficient of each mutation observed at frequencies above the sequencing error threshold. We thus utilized WFABC (Foll *et al.* 2014; Foll *et al.* 2015) to identify positively selected candidate mutations (see Materials & Methods). All considered trajectories are displayed in Supplementary Figure 5. Supplementary Table 2 contains a list of all identified candidates. Of note, candidate mutations were identified by WFABC under the assumption of independence between sites, an assumption likely violated in IAV. Hence, each candidate may be the subject of either direct or linked selection. However, WFABC was successful in identifying resistance mutations against oseltamivir (Foll *et al.* 2014), several of which have been functionally validated; thus, it was a valuable tool for identifying a set of promising candidates for favipiravir.

Overall, WFABC identified 64 positively selected candidate mutations, of which 24 were shared between at least two populations. The allele-frequency trajectories of all candidate mutations across all populations are plotted in Supplementary Figure 6. Strikingly, the strongest signal of selective sweeps (see also Supplementary Figure 5) and the highest number of candidate mutations (*n*=21, across passages 3-19) were seen in the constA population, which also showed signs of resistance evolution based on recovering growth rates, as detailed in the previous subsection (see Figure 3D); these candidates also had the highest inferred selection coefficients (see Supplementary Table 2). Of the 21 candidate mutations, 10 are non-synonymous and the majority (*n*=8) are at genes localized to the subunits of the viral RdRp, which could be expected given that favipiravir is proposed to inhibit the viral RdRp (Furuta *et al.* 2005; Jin *et al.* 2013). The phenotype of most of the candidate mutations is unknown.

A well-characterized candidate within this group, NP S9T, disrupts a phosphorylation site and alters nuclear-cytoplasmic shuttling of the nucleoprotein (NP) (Hutchinson *et al.* 2012; Zheng *et al.* 2015). Phosphorylation of NP S9 prevents nuclear import, and mutations at this site thus lead to an accumulation of NP within the nucleus (Zheng *et al.* 2015), the location of viral RNA replication. Altering the levels of NP in the nucleus could increase rates of viral RNA replication (Portela and Digard 2002), thereby counteracting the inhibitory effects of favipiravir, though this mechanism needs to be formally tested. Interestingly, NP S9 is perfectly conserved in H1N1 isolates, and thus the allele may be an unlikely resistance mutation in natural isolates.

Additional candidate mutations were identified in the constB population (n=6). One of the candidates, NP D101N, was previously identified in an experimental screen for alleles conferring resistance to the drug ribavirin (Cheung *et al.* 2014). Like favipiravir, ribavirin increases the mutation rate of the IAV genome (Cheung *et al.* 2014). Interestingly, follow-up assays could not confirm the action of NP D101N as a resistance mutation (Cheung *et al.* 2014), suggesting that either NP D101 is a product of cell-culture adaptation, or perhaps contributes to resistance only in interaction with other mutations. Analysis of our previously published data on serial passaging of IAV in MDCK cells (Foll *et al.* 2014; Renzette *et al.* 2014) showed fixation of NP D101N in one of the two trajectories where oseltamivir-resistant influenza virus evolved, which suggests that NP D101 indeed emerges with adaptation to MDCK cells. Of note, the PB1 V43I and the PB1 D27N mutations, known to affect polymerase fidelity and confer resistance to ribavirin in other studies (Cheung *et al.* 2014, Binh *et al.* 2014), is not detected in our favipiravir experiments.

In both the favi1 and favi2 populations, few (*n=2* for favi1 passages 3-14; *n*=3 for favi2 passages 3-10) candidate mutations were identified before the population was driven to extinction. The mutations in this group were distributed across the genome, suggestive of the absence of a strong selective sweep. In the control and withdrawal populations, the identified candidates (listed in Supplementary Table 2) were largely shared between populations, indicating potential adaptations to the MDCK environment. Indeed, many of the candidate mutations across populations have previously been described in the literature associated with mammalian cell culture adaptation (e.g. PB2 G590S (Mehle and Doudna 2009; Poole *et al.* 2014) and HA2 D112N (Foll *et al.* 2014).

### Potential drivers of adaptation in constA

To identify groups of mutations indicative of genetic hitchhiking and/or joint selection in the constA environment, we performed a hierarchical clustering analysis based on the squared Euclidean distance between allele-frequency trajectories (see Materials & Methods). The result of this analysis is illustrated in Figure 4. We chose a dissimilarity threshold of 0.95, which results in six clusters (see Figure 4A).

The mutations in cluster 1 (see Figure 4B) begin to increase in frequency at passage 9, though come to a halt (potentially due to interference) around passage 13, which also coincides with the increasing frequency of cluster 5. Two non-synonymous mutations (indicated with an asterisk in Figure 4A) are the likely targets of direct selection (i.e., drivers) in this cluster: HA S220P is a mammalian cell adaptation that has been observed in MDCK cells, ferrets and humans (Smirnov *et al.* 2000; Ding *et al.* 2010; Imai *et al.* 2012); the same mutation is also identified as a candidate in the favi2 population. PB2 G590S is a temperature-sensitive polymerase mutation that has been frequently observed upon switching from propagation of the virus in chicken eggs to mammalian cells (Mehle and Doudna 2009; Poole *et al.* 2014). Hence, this cluster likely represents an adaptation to the cell culture.

Cluster 2 contains only a single, non-synonymous mutation in the polymerase subunit PB2 that has not been characterized previously. The frequency of this mutation decreased towards the end of the experiment, which may be indicative of linked rather than direct selection, for example from clusters 1 or 3.

Cluster 3 was the earliest to begin increasing in frequency in the population, and several of the mutations contained in this cluster were present early in the experiment (during the phase of increasing drug concentration; see Figure 4D). The best-characterized candidate and potential driver of this cluster is NP S9T (discussed above). However, an additional potentially interesting candidate for further study is PA R204R/PA-X D204G, which represents a non-synonymous mutation in the recently discovered second open reading frame of segment 3, and whose protein product has been reported to be involved in virulence of IAV (Jagger *et al.* 2012).

Cluster 4 is represented by two mutations showing highly uncharacteristic allele-frequency trajectories (see Figure 4E). Given their long persistence at intermediate frequencies, they are unlikely to be driver mutations.

Cluster 5 contains the highest number of mutations, with four being tightly linked in the polymerase subunit PA. Most of the involved mutations are synonymous, making them unlikely driver candidates. Two non-synonymous mutations in PA, E31G and E56G, have not been previously characterized. However, the third non-synonymous mutation in this cluster, PB2 K718E mutates a residue important in PB2 binding to various importin α isoforms (α1, α3 and α7), and thus is critical in altering the kinetics of PB2 nuclear importation (Pumroy *et al.* 2015). Combining this putative phenotype with that of the NP S9T allele of cluster 3 suggests a model in which adaptation to favipiravir is associated with alteration of the sub-cellular localization of viral RdRp components rather than changes in viral RdRp enzymatic activity or drug binding. This model is tentative, though, and warrants further investigation.

Finally, cluster 6 contains two mutations, one of which is synonymous. The other mutation is outside the protein-coding domain of the polymerase subunit PB2 and is of unknown function.

In summary, the following observations emerged from the cluster analysis: first, larger clusters (suggestive of stronger selective sweeps) contained the most compelling candidate mutations for adaptation and resistance evolution. Second, we were able to reconstruct a hypothesized history of adaptation: an early selective sweep of MDCK cell adaptation (cluster 1) interferes with a later sweep of a resistance mutation (cluster 5). However, a high mutation rate and co-infection (and, hence, reassortment of segments) appears to enable the combination of multiple beneficial mutations on the same background, which leads to rapid adaptation in this population. It is unclear whether the staggered sweeps observed here would be possible in isolation or whether the effects of later mutations are indeed dependent on the presence of earlier mutations. However, our results provide an excellent means of identifying candidates for functional testing.

**Figure 4:**
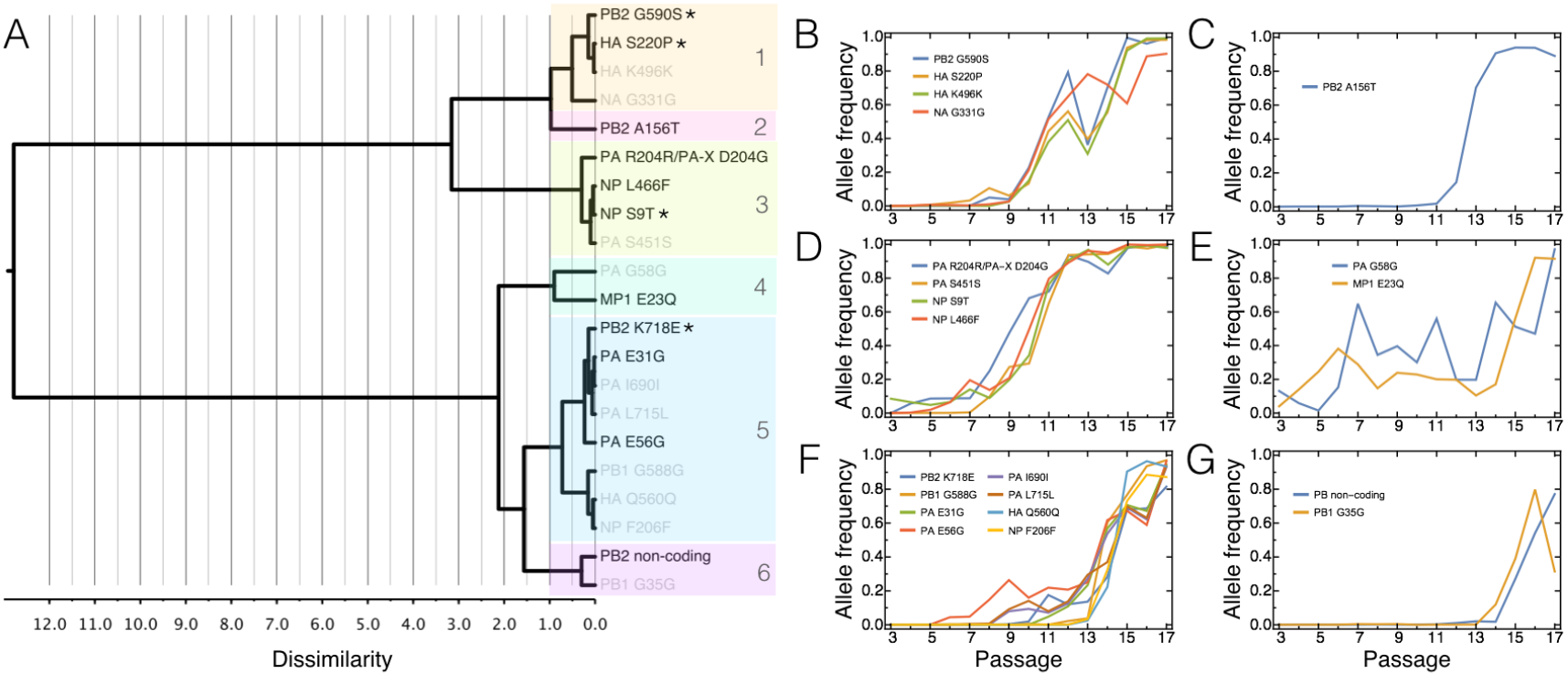
Clustering of WFABC candidate mutations in the constA environment. Panel A: Tree showing the (dis-)similarity between allele-frequency trajectories of beneficial candidates as an indicator of either hitchhiking effects or joint selection. Asterisks indicate the mutations that are discussed in the main text as potential driver mutations. Panel B-G: Allele-frequency trajectories of the mutations in each cluster.

### Assessment of changes in the selection pressure

We also apply a newly developed extension of WFABC to test for severely changing selection coefficient *s* or population size *N_e_* during the course of the experiment. CP-WFABC (“change-point”-WFABC (Shim *et al.* 2016); see Materials and Methods) can also identify the change point and magnitude of the product of *N_e_* and *s* before and after the change, and tests a model with changing parameters against the null model of a single fixed selection coefficient and population size. As visualized in Supplementary Figure 7, the results of CP-WFABC were consistent with the rapid population extinction in favi1. The population collapse was accompanied by a steep decrease in the effective population size, which produces hundreds of trajectories that are not consistent with a classical constant-environment model. In contrast, only few and relatively uniformly distributed change points are identified in the control populations, consistent with our expectation in a constant environment.

### Favipiravir-induced mutational meltdown

We observed successful extinction of the viral population within <150 generations in both replicates under increasing drug concentrations. The observed pattern is stunningly similar to the dynamics of mutational meltdown described in Lynch *et al.* (1993): shortly after a new asexual lineage is created, individuals accumulate mutations almost linearly until reaching a transition point at which the population size collapses, producing a sharp increase in mutation accumulation. In Figure 5, the accumulation of mutations per individual (see Materials & Methods) in the favi1 population (blue dots) is overlaid on a reproduction of Figure 1 of Lynch *et al.* (1993) (blue line). Although we lack information on the carrying capacity of the population and the census population size (shown as gray dashed line following Lynch *et al.* (1993)), the observed changes in the effective population size (gray dots) are consistent with this model: over time, the effective population size decreases gradually (due to Muller’ s ratchet) but slowly, until it collapses at the transition point. Notably, this transition point was also indicated by CP-WFABC as a major change in the selection pressure. The same pattern was observed for the favi2 population, and all populations showed the linear mutation accumulation expected under Muller’ s ratchet (visualized in Supplementary Figure 8). Because mutation accumulation in the model is proportional to the mutation rate, comparing the slopes provides additional confirmation of the increase in mutation rate observed under favipiravir treatment (see Supplementary Figure 8).

**Figure 5.**
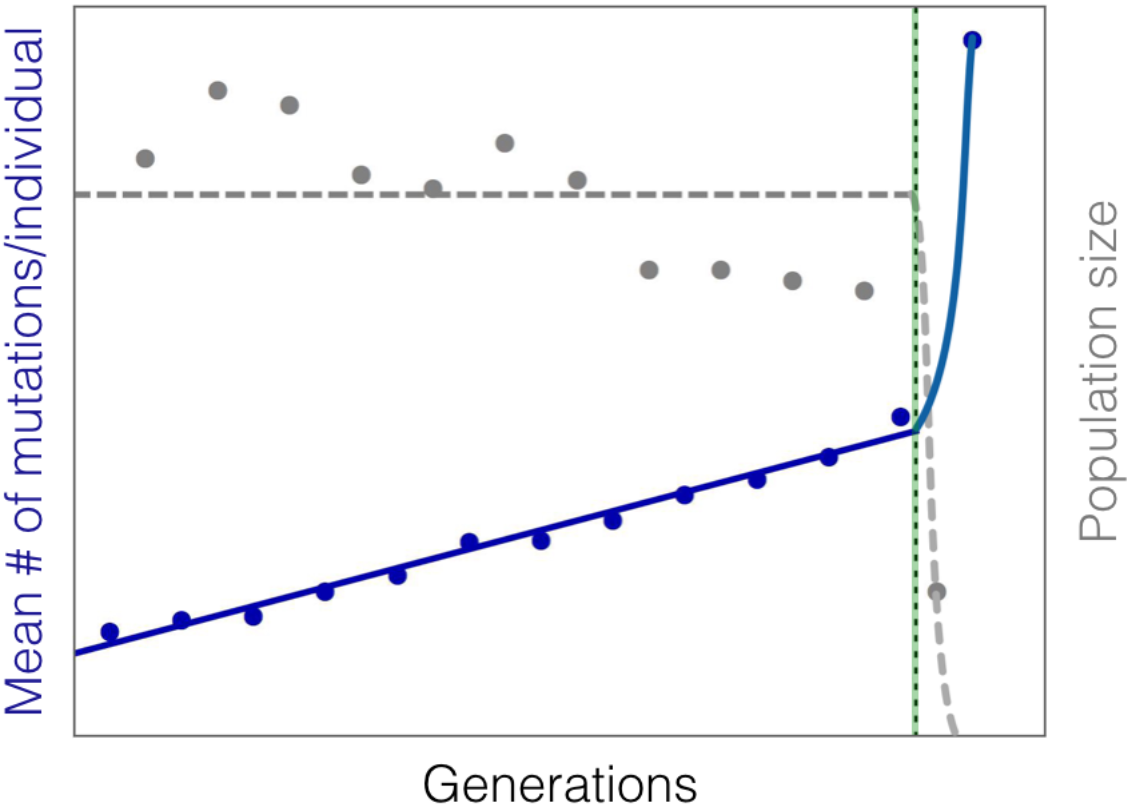
Qualitative comparison of accumulation of mutations in the favi1 population (blue dots) with pattern of mutation accumulation redrawn from Figure 1 of Lynch *et al.* (1993) as the solid blue line, representing mutational meltdown. The horizontal gray dashed line represents the census population size in the original model, which is here overlaid by our *N_e_* estimates between passages (gray dots). The dashed vertical line in green indicates the transition to the meltdown phase.

## CONCLUSION

In this study, we examined a novel class of drug treatment that acts to increase viral mutation rates. Although an increased mutation rate may allow for a more rapid appearance of beneficial mutations in the population (e.g., Cirz and Romesberg 2007), the underlying notion is that the comparatively much greater input rate of deleterious mutations should lead to population extinction (i.e., within-host extinction of the virus) and prevent any rescue mutations from emerging. This expectation is supported by a large body of classical studies in evolutionary theory that have described the processes of mutational meltdown (e.g., Lynch *et al.* 1993), lethal mutagenesis (e.g., Bull *et al.* 2007; Martin and Gandon 2010; Arias *et al.* 2014) and error catastrophe (e.g., Biebricher and Eigen 2005).

By comparing multiple replicates of populations grown in the absence of favipiravir treatment, at constant concentrations of the drug, and at escalating concentrations, we quantified the respective effects on underlying mutation rates and described the resulting evolutionary processes. We demonstrated that all populations treated with increasing concentrations of favipiravir were characterized by increased mutation rates, decreasing effective population sizes through time, no observed rescue mutations, and ultimate extinction in all population replicates (see Table 1). This pattern is in sharp contrast to populations treated with a constant concentration of favipiravir, which maintain constant (but reduced) effective population sizes, show signs of selective sweeps and, in one replicate, a striking recovery of the population growth rate - suggesting that lower concentration conditions may indeed allow for the virus to persist and potentially even develop resistance. The contrast with populations grown in the presence of oseltamivir is also noteworthy, for which single-mutational-step resistance mutations rapidly arose in all treatment populations allowing for a complete evolutionary rescue (Foll *et al.* 2014).

**Table 1.**
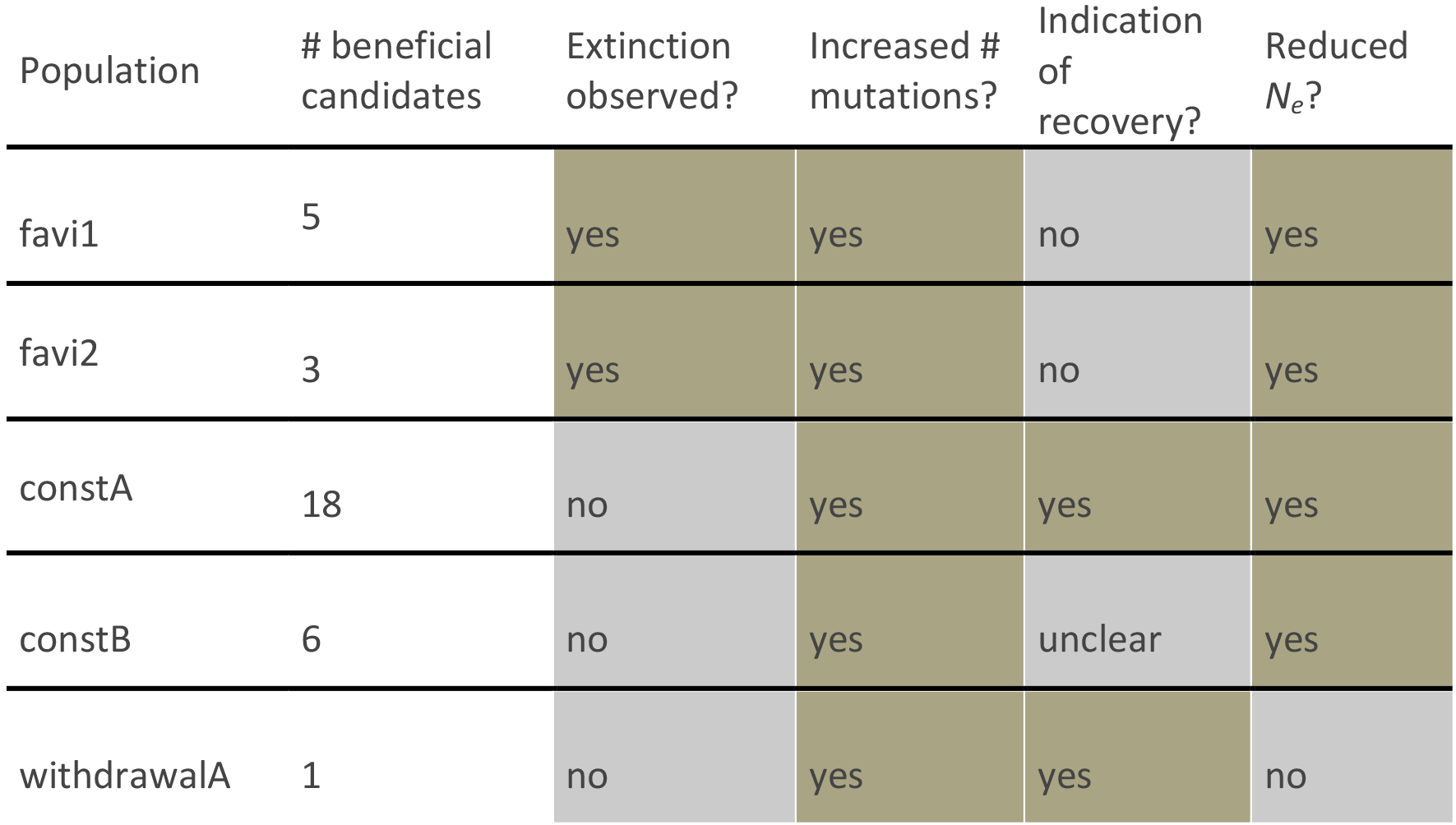
A summary of observations across data sets

| Population | # beneficial candidates | Extinction observed? | Increased # mutations? | Indication of recovery? | Reduced $N_e$? |
| --- | --- | --- | --- | --- | --- |
| favi1 | 5 | yes | yes | no | yes |
| favi2 | 3 | yes | yes | no | yes |
| constA | 18 | no | yes | yes | yes |
| constB | 6 | no | yes | unclear | yes |
| withdrawalA | 1 | no | yes | yes | no |

As the ultimate source of variation, mutational effects and rates have remained a persistent subject in evolutionary theory. In 1930, R.A. Fisher (1930) argued that an intermediate mutation rate is optimal for organisms to ensure a steady input of beneficial mutations while avoiding the detrimental accumulation of deleterious mutations. His arguments were later formalized in several evolutionary concepts including Muller’s ratchet (Muller 1964; Felsenstein 1974), mutational meltdown (Lynch *et al.*, 1990), lethal mutagenesis (Bull *et al.*, 2007), background selection (Charlesworth *et al.* 1993; Charlesworth 2012), background trapping (Johnson and Barton 2002), Hill-Robertson interference (Hill and Robertson 1966; McVean 2000), and, in a more biophysically inspired framework, quasi-species theory (Eigen 1971; Biebricher and Eigen 2005; but see Wilke 2005). Whereas these models generally predict an eventually detrimental effect of increasing the mutation rate, instances of rapid resistance evolution against mutation-rate increasing treatment (Pfeiffer and Kirkegaard 2003) and a general escape from extinction (Springman *et al.* 2010) have been previously reported.

Furthermore, so-called mutator genotypes are frequently observed when bacteria are exposed to novel environments, where an increased mutation rate may facilitate adaptation, particularly over short time scales (Taddei *et al.* 1997; Ram and Hadany 2012). Therefore, efforts are made to develop mutation-inhibition treatments to prevent antibiotic resistance evolution in bacterial pathogens (Cirz and Romesberg 2007). Conversely, as demonstrated here and as previously argued theoretically (e.g., Martin and Gandon 2010), increasing mutation rates indeed also represent a potential treatment strategy.

Thus, the precise relevance of this information for the study of virus evolution and the development of improved treatment strategies requires further examination. First, the correspondence of the observed patterns with classical theory demonstrates the predictive value of population-genetic models. In the model of Lynch *et al.* (1993), extinction time is estimated based on the mutation rate, the carrying capacity, the rate of reproduction, and the selection coefficient. Whereas the relationship between the reproductive rate and the carrying capacity and extinction time are relatively simple, we here present a novel finding of the (deleterious) selection coefficient having a non-linear relationship with extinction time, a time that is minimized under intermediate selection coefficients. This has important implications for the evolution of the virus: if changing the environment (e.g., drug pressure) changes the distribution of fitness effects of new mutations, this can result either in shorter or longer extinction times. Changing this distribution also alters the relevance of the discussed evolutionary mechanisms (e.g., Muller’ s ratchet, background selection, WSHRI). Essentially, minimizing the expected extinction time optimizes drug efficacy and decreases the risk of resistance evolution. By combining our emerging knowledge of the underlying distributions of fitness effects of new mutations with classical theory, we may be able to develop better predictions regarding the efficacy of both single and combination drug therapies.

Second, we observe that the number of accumulated mutations per individual in the passage immediately prior to extinction was almost twice as large in the favi1 as compared with the favi2 population. This may be partly explained by differences in the experimental setup (see Materials & Methods), but considering the similar effective population sizes it more likely provides evidence for the inherent stochasticity of the extinction process (Lynch *et al.* 1993; Martin and Gandon 2010; Wylie and Shakhnovich 2012). Hence, it supports the synergism between stochastic and deterministic drivers of extinction proposed in the theory of mutational meltdown (Lynch *et al.*, 1993) rather than error catastrophe, which proposes extinction due to the inability of the population to contain information upon crossing a (sharp) error threshold.

Thus, this work is an important empirical insight into the widely theorized models discussed above. Further, by experimentally controlling the demographic dynamics of the population as well as the imposed selective pressures, we avoid many of the commonly confounding effects encountered in attempts to quantify these processes. In addition, the results of this evolutionary study hold great clinical importance, as they validate the notion of inducing mutational meltdown as a viable viral treatment strategy.

Importantly, our results indicate the great influence of dosage, and present the first evidence to date for viral adaptation to favipiravir treatment. Encouragingly however, under high concentration environments rescue mutations are not observed, ultimately resulting in population extinction. The mechanism of action of favipiravir is hypothesized to be of relevance across RNA viruses, and the results presented here thus warrant future comparative studies in, for example, Ebola virus and West Nile virus populations. With regards to IAV specifically, these results are encouraging for the future promise of improved treatment strategies to help minimize the great public health costs of this virus.

## ACKNOWLEDGMENTS

We are grateful to Kristen Irwin, Matt Jones, Stefan Laurent, Yoav Ram, Melanie Trombly, Severine Vuilleumier, and Alex Wong for helpful comments on the manuscript.

